# Dataset-adaptive minimizer order reduces memory usage in *k*-mer counting

**DOI:** 10.1101/2021.12.02.470910

**Authors:** Dan Flomin, David Pellow, Ron Shamir

**Affiliations:** Blavatnik School of Computer Science, Tel-Aviv University, Tel-Aviv, Israel

## Abstract

The rapid, continuous growth of deep sequencing experiments requires development and improvement of many bioinformatics applications for analysis of large sequencing datasets, including *k*-mer counting and assembly. Several applications reduce RAM usage by binning sequences. Binning is done by employing minimizer schemes, which rely on a specific order of the minimizers. It has been demonstrated that the choice of the order has a major impact on the performance of the applications. Here we introduce a method for tailoring the order to the dataset. Our method repeatedly samples the dataset and modifies the order so as to flatten the *k*-mer load distribution across minimizers. We integrated our method into Gerbil, a state-of-the-art memory efficient *k*-mer counter, and were able to reduce its memory footprint by 30% - 50% for large *k*, with only minor increase in runtime. Our tests also showed that the orders produced by our method produced superior results when transferred across datasets from the same species, with little or no order change. This enables memory reduction with essentially no increase in runtime.

## 1 Introduction

High-throughput sequencing (HTS) has enabled rapid progress in biological and clinical research through efficient and cheap sequencing of large genomic and transcriptomic samples. Analyzing the sequences from large HTS experiments presents computational and algorithmic challenges due to the high volume of data and the fragmented sequences generated. *k*-mer counters are a fundamental building block in some of the most basic tasks in analyzing HTS data, including genome assembly [7], repeat detection, and multiple sequence alignment.

Over the past few years, several *k*-mer counters were developed. A main paradigm to speed-up and reduce memory footprint of *k*-mer counting is binning [3, 4, 5, 11, 12]. Such methods partition the entire dataset into several bins, and then process each bin independently (possibly in parallel) before combining the results into the final output. Gerbil [5] and KMC3 [9] are popular *k*-mer counting tools that use binning, but differ in their counting approach: KMC3 sorts each bin, while Gerbil employs hash tables. BCALM2 [3] is a genome assembly tool that uses a *k*-mer counter similar to KMC2, an earlier version of KMC3, as a pre-process to its assembly phase.

Minimizers [17] have been broadly used to speed up sequence analysis algorithms and to reduce disk space and memory. Given integers *k* and *m*, the *minimizer* of a *k*-long sequence (*k*-mer) is the smallest among the *k* − *m* + 1 contiguous *m*-mers in it, where the smallest is determined based on a predefined order (e.g., lexicographic or random). For a longer sequence, all *k*-long contiguous substrings are scanned and the minimizer is selected in each one (Figure 1).

**Figure 1:**
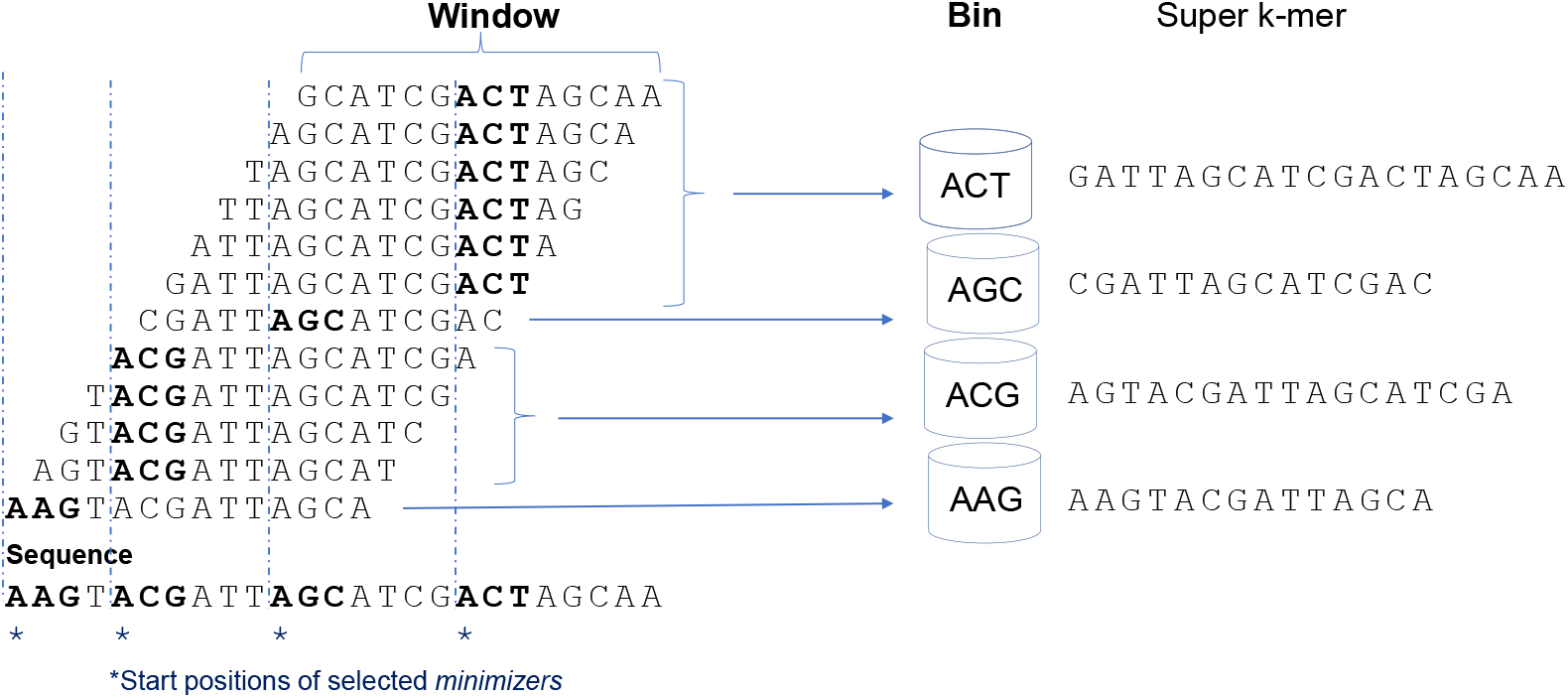
Illustration of a minimizer scheme and binning. Here *k* = 12 and *m* = 3. The input sequence is broken into windows of length *k*, and in each window the *m*-long minimizer according to the lexicographic order (shown in bold) is selected. The *k*-mer is assigned to the partition whose label is the minimizer. Consecutive windows tend to select the same minimizer, and the concatenation of the consecutive windows forms the super-*k*-mer that is stored in the bin.

In sequencing applications, the *k*-mers in the dataset are partitioned using their minimizers as the key, i.e., each *k*-mer is assigned to the partition corresponding to its minimizer. Since the same *m*-mer is often selected from overlapping windows, partitioning with the minimizer as key helps compress the data by representing several overlapping *k*-mers with the same minimizer as a longer super-*k*-mer in the minimizer’s partition [12]. Often, partitions are grouped into a smaller number of bins.

Minimizer schemes have their limitations. Certain minimizers tend to appear much more than others in biological data, leading to highly unbalanced bin sizes. For example, the bin containing the minimizer *AA*…*AA* tends to be very large when using a lexicographic order. Several applications tried to overcome this problem by proposing alternative orders. KMC2 [4] proposed the signature order, which tries to avoid certain *m*-mers such as *AA*..*A*. The Minimap2 read mapper [10] uses a random order of the minimizers. Defining an order based on minimizers taken from a universal hitting set (UHS) was demonstrated to reduce density and increase mean distance between selected minimizer positions [14]. That order was shown to reduce memory in genome assembly [1].

Some methods introduced minimizer orders based on the statistics of the particular sequences of interest. Chikhi et al. [2] showed that ordering *m*-mers by increasing frequency in the data dramatically reduced memory usage in assembly graph construction compared to both lexicographic and random orders. In [16], the UHS order was combined with the frequency based order to balance bin sizes and lower memory in distributed *k*-mer counting. In the Winnowmap read mapper [8] the most frequent *m*-mers in the dataset are moved to appear later in the order.

The number of distinct *k*-mers associated with a minimizer in a particular dataset is known as the *load* of the minimizer [1]. In applications that partition the data based on minimizers, hash multiple partitions into bins, and process each bin separately, it is desirable to have bins with smaller load, in order to ensure low RAM usage, as has been argued by [12].

In this study we introduce a new approach for adapting the minimizer order to the target sequence data. The method, called AdaOrder (Adaptive Order), iteratively updates the minimizer order based on an estimate of the minimizer loads in the data in order to reduce the maximum load. We demonstrated its ability to lower the maximum load compared to all predefined orders, except the frequency based order.

We integrated our new order into Gerbil [5], a leading *k*-mer counter, which uses signature order, and was consistently the most memory-efficient in a recent benchmark [13]. In tests on several datasets, our implementation, called DGerbil, achieved a reduction of up to 50% in memory usage for large values of *k*, with only slightly higher running times compared to Gerbil. Our tests also showed that the orders produced by our method produced superior results when transferred across datasets from the same species, with little or no order change. This enables memory reduction with essentially no increase in runtime. We also implemented Gerbil with the frequency based order [2], and showed that DGerbil required significantly less memory.

The code of AdaOrder and DGerbil, as well as orders produced for the studied datasets and species, are publicly available at github.com/Shamir-Lab/AdaOrder.

## 2 Definitions and Background

### 2.1 Basic definitions

#### Reads and *k*-mers

A *read* is a string over the DNA alphabet Σ = {*A, C, G, T*}. An *ℓ*-mer is an *ℓ*-long string over Σ. For an *ℓ*-mer *s* with *k* ≤ *ℓ* and 0 ≤ *i* ≤ *ℓ* − *k* we define *s*(*i*) to be the *i*th *k*-mer in *s*. By convention, positions in *ℓ*-mers start at zero.

#### Natural *m*-mer mapping

We treat each *m*-mer as a number in base 4, with the following DNA base encoding: *A* = 0, *C* = 1, *G* = 2, *T* = 3. For example, with *m* = 4, *AAAA* = (0000)_4_ = 0, *TTTT* = (3333)_4_ = 255, *ACGT* = (0123)_4_ = 27. The natural mapping is denoted as *Natural* : Σ^*m*^ → {0, …, 4^*m*^ − 1}. We refer to *m*-mers and values in {0, …, 4^*m*^ − 1} interchangeably.

#### Reverse complement and canonical form

Given an *m*-mer *x*, its *reverse complement* is constructed by reversing the order of the bases in *x* and taking their complements. The complements of bases *A, C, G, T*, are *T, G, C, A* respectively. Given an *m*-mer *x*, 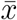 denotes its reverse complement. For example, 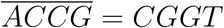.

The *canonical form* of an *m*-mer *x* is the smaller of *x* and 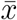 with respect to the natural mapping. We denote *x*’s canonical form by *Canonical*(*x*). For example, *Canonical*(*CGGT*) = *ACCG*.

We refer to the set of all canonical *m*-mers as *c*_*m*_ = {*Canonical*(*x*)|*x* ∈ Σ^*m*^}.

#### *m*-mer order

An *order o* on Σ^*m*^ is a function *o* : Σ^*m*^ → ℝ. An order *o* can be *normalized* to *o*_*norm*_ : Σ^*m*^ → {0, 1, …, |Σ|^*m*^ − 1} (breaking ties consistently). Here and throughout, we treat *x* and 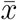 as equivalent. In this notation *o*(*x*) ≡ *o*(*Canonical*(*x*)) for any *x* in Σ^*m*^ and so 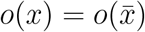 and *o* : *c*_*m*_ → ℝ.

#### Minimizers

A *minimizer* of a sequence *s* with respect to order *o* is the smallest *m*-long contiguous substring *z* in *s* according to *o*. We also call *z* the *o*-minimizer *m*-mer in *s*.

Within a sequence, *m*-mer *x* is smaller than *m*-mer *y* according to an order *o* if: (i) *o*(*x*) *< o*(*y*); or (ii) *o*(*x*) = *o*(*y*) and *Natural*(*x*) *< Natural*(*y*); or (iii) *o*(*x*) = *o*(*y*) and *Natural*(*x*) = *Natural*(*y*) and *x* occurs to the left of *y* in *s*. In other words, we break ties according to *Natural*, and if needed also by choosing the leftmost appearance of the least *m*-mer.

A *minimizer scheme* is a function *f*_*o,m,k*_ : Σ^*k*^ → [0 : *k* − *m*] that selects the start position of the *o*-minimizer *m*-mer in every sequence of length *k* (**Figure 1**).

#### Lexicographic order

We define the lexicographic order to be: *o*_*lexico*_(*x*) = *Natural*(*x*)

#### Signature order

The following *signature order* was proposed in [4] as an alternative to the lexicographic order. Each *m*-mer that either starts with AAA or ACA or contains AA as a substring is called *bad*. The rest of the *m*-mers are called *good*. The good *m*-mers are ordered lexicographically, and all of them are defined to be smaller than the bad *m*-mers. The bad *m*-mers are not ordered and are all mapped to the same value.

We define a variant of the signature order as follows:

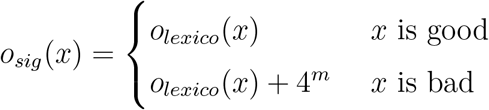

Note that unlike in [4] we do order the bad *m*-mers.

#### Random order

We define a pseudo-random order by performing a xor operation between the binary representations of a random integer mask *α* and of the lexicographic order of *m*-mers. We denote ⊕ as the bitwise xor operation. The random order is formally defined as follows:

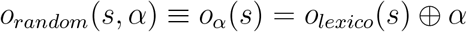

The xor operation is a bijection, and therefore if *s*≠ *s*′ then *o*_*α*_(*s*) ≠ *o*_*α*_(*s*′).

#### Frequency order

The *frequency order* [2] is a minimizer order based on statistics collected from a specific dataset. It orders the *m*-mers in increasing frequency order according to the dataset. For our analysis we defined it as follows: Given a dataset *D*, we count *m*-mer appearances in it, and set

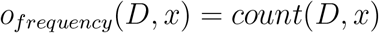

where *count* is the number of times *x* appeared in *D*.

#### Partitions, partition size, bins and super-*k*-mers

In *minimizer partitioning* each *k*-mer in the input dataset is assigned to a partition (e.g., a file on disk) according to its minimizer. Each partition corresponds to only one minimizer and vice versa. When the number of partitions is too large, partitions are often combined into a smaller number *B* of groups called *bins*, where each bin contains the *k*-mers of several minimizers, and each minimizer corresponds to only one bin. The *size* of a partition or a bin is the total number of characters in all *k*-mers in it.

Several consecutive *k*-mers that share the same minimizer can be compressed together. A *super-k-mer* [12] is the longest substring for which all *k*-mers have the same minimizer. **Figure 1** shows an example of partitions and super-*k*-mers.

### 2.2 Quality Criteria for minimizer schemes

Following [1, 16], we define quality criteria for a minimizer order on a dataset. They are meant to indicate how evenly the *k*-mers of a dataset are distributed across the minimizers.

#### Load

Given a minimizer scheme *f*_*o,m,k*_ and a dataset *D*, the *load* of *x* ∈ *c*_*m*_ is the number of distinct *k*-mers in *D* for which *x* is the minimizer [1]. We denote it by *l*(*x, D*). Similarly, when binning is used, we define the bin’s load to be the number of distinct *k*-mers in it.

We also define the *relative load* of *x* ∈ *c*_*m*_ over dataset *D* to be *x*’s load divided by the total number of distinct *k*-mers in the dataset. We denote it by *r*(*x, D*).

The *maximum* (relative) load is the (relative) load of the minimizer with the highest (relative) load. Maximum (relative) load is defined analogously for bins.

#### Unevenness

Unevenness aims to describe how balanced a minimizer scheme is over a particular dataset.

Given a minimizer scheme *f*_*o,m,k*_ and a dataset *D*, we define its unevenness to be:

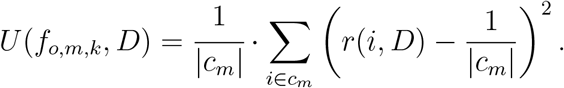

Recall that |*c*_*m*_| is the number of possible *m*-long minimizers, and thus the unevenness is the mean squared difference from the uniform distribution of *k*-mers across minimizers.

### 2.3 The Gerbil *k*-mer counter

Gerbil [5] is a memory efficient *k*-mer counter. It bins *k*-mers according to their minimizers, using the signature order in its minimizer scheme, and merges adjacent *k*-mers sharing the same minimizer into super-*k*-mers. Then the *k*-mers in each bin are counted independently using a hash table.

### 2.4 Mapping minimizers to bins in Gerbil and KMC3

*k*-mer counters use a small, fixed number of bins (e.g., 512 in Gerbil) and assign multiple partitions to the same bin. The mapping of minimizers into bins could be performed randomly (e.g., using a random hash), but many *k*-mer counters use heuristics that aim to better balance the loads of the bins. We note that achieving an optimal distribution of minimizers to bins is equivalent to the NP-Hard problem of optimal identical machines scheduling with minimum makespan [6]. There exist complex heuristics and polynomial time approximation schemes for this problem [18], however, in practice *k*-mer counting tools use simple heuristics without guarantees, in the interest of runtime and simplicity.

Gerbil tries to group together minimizers that appear earlier in the order (and thus are intuitively likely to have higher load) with those that appear later. Specifically, each minimizer is added to a new bin in increasing signature order. The order of the bins to which minimizers are added is reversed each time a minimizer has been added to every bin.

KMC3 [9] samples a portion of the dataset in order to estimate the partition size of each minimizer. After collecting the statistics it adds 1000 to each partition size and then sorts the minimizers by their sampled counts from the largest to the smallest. Minimizers are mapped to the current bin until its size exceeds the average remaining bin size, and then a new bin is opened, and the process continues in that bin. An outline of the process is described in **Supplementary Algorithm S1**.

## 3 Methods

We developed an algorithm called AdaOrder that constructs a minimizer order with low maximum load for a given dataset, based on statistics collected on the dataset. We also implemented a sampling process that estimates the total size of each partition. These estimates are used to determine an efficient mapping of minimizers to bins, similar to KMC3. The process is schematically as follows: (i) Compute a minimizer order using AdaOrder; (ii) Estimate the minimizer partition sizes by sampling; (iii) Map minimizers to bins using the estimates from (ii). We integrated this process into Gerbil and call the modified algorithm DGerbil.

### 3.1 AdaOrder

AdaOrder is a heuristic that aims to produce a minimizer order with low maximum minimizer load in a given dataset. AdaOrder starts with the scheme 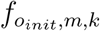 and works in *R* rounds. In each round it samples *N k*-mers from the dataset in order to capture a minimizer with a large load. A single round proceeds as follows: (i) Iterate over the dataset’s reads sequentially; (ii) For each read scan through all of its *k*-mers and identify minimizers; (iii) Compute the load of each minimizer according to the sampled *k*-mers; (iv) After sampling at least *N k*-mers identify the minimizer *µ* with the highest sample load. In case of ties, choose the lexicographically smallest minimizer. Alter the current order by increasing the order of *µ* by *p* · 4^*m*^, where *p* is a penalty factor. This makes *µ* less likely to be chosen as a minimizer, thus lowering its load. **Algorithm 1** describes AdaOrder.

#### Algorithm 1 AdaOrder

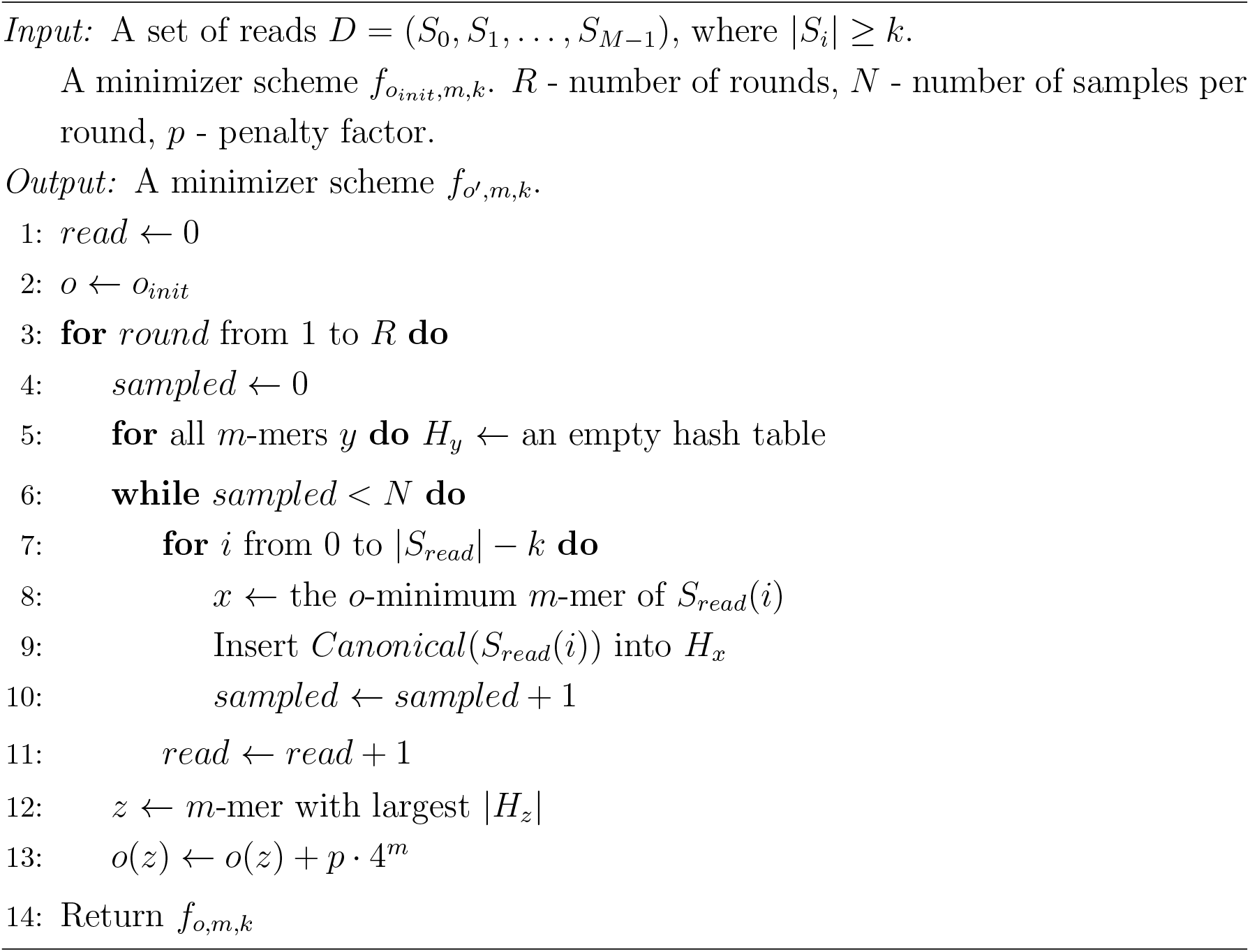

### 3.2 Mapping minimizers to bins

We first estimate each partition size through sampling (**Supplementary Algorithm S2**), and then map minimizers to bins using these estimates. The bin mapping algorithm is shown in **Algorithm 2**; it heavily relies on KMC3’s implementation.

### 3.3 DGerbil

DGerbil is based on the memory efficient *k*-mer counter Gerbil. It integrates AdaOrder initialized with signature order and our bin mapping algorithm into Gerbil. The use of AdaOrder aims to lower the maximum bin load, in order to lower the RAM usage. The algorithm for DGerbil is shown in **Algorithm 3**. Gerbil’s binning method and counting method are described in **Section 2.3**.

#### Algorithm 2 BinMapping

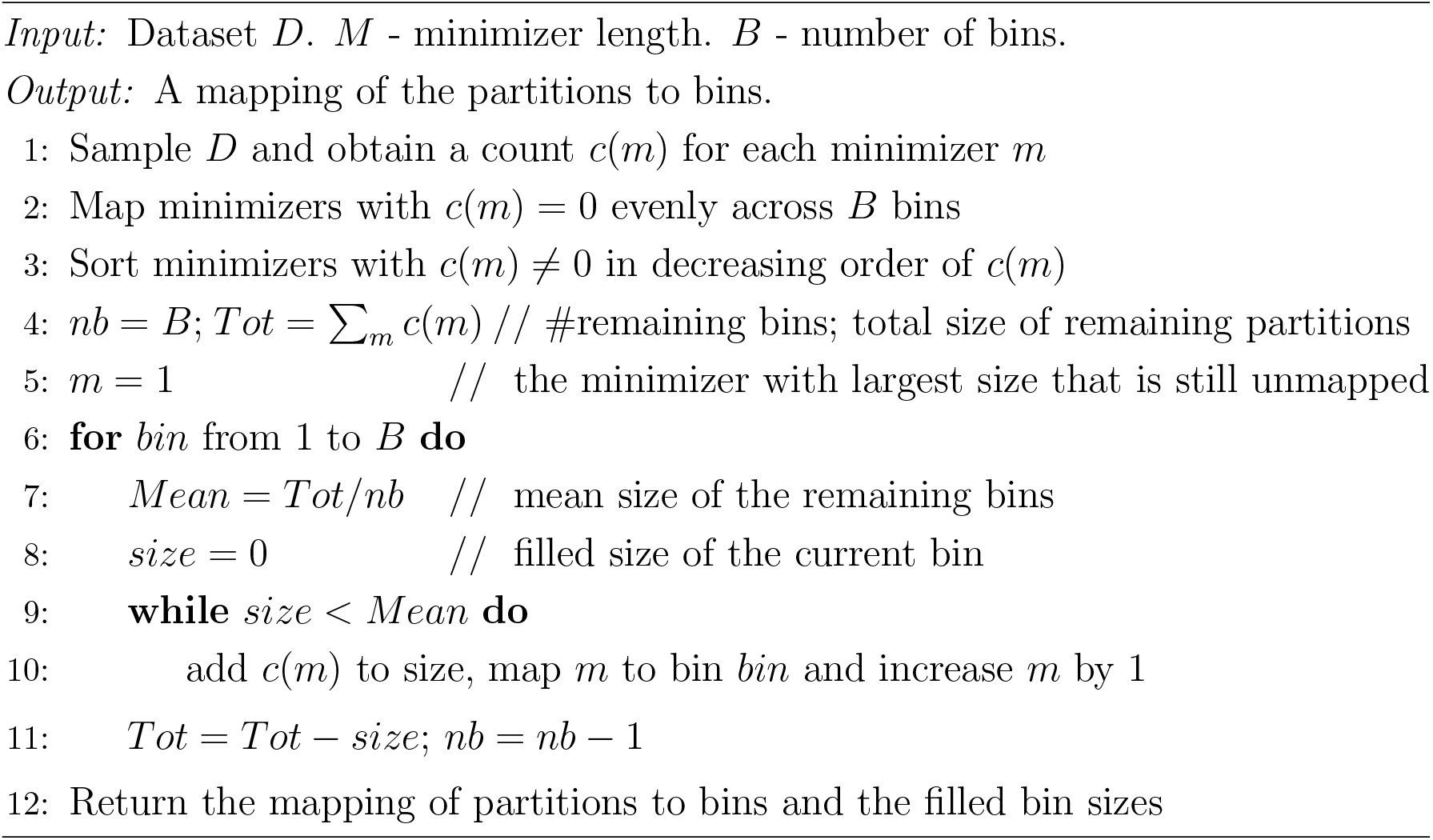

### 3.4 FGerbil

We implemented a variant of Gerbil that uses frequency order, called FGerbil. It first counts how many times each *m*-mer appeared in the dataset, and orders *m*-mers by their frequency. It then uses the resulting frequency order within Gerbil, and performs bin mapping as DGerbil does.

### 3.5 Source code

The methods and code base to run AdaOrder and DGerbil are available at github.com/Shamir-Lab/AdaOrder. AdaOrder is coded in Java, while DGerbil is a modification of Gerbil’s C++ code.

In our implementation of AdaOrder some optimizations were introduced to **Algorithm 1** for a faster running time (e.g., keeping the previous minimizer and the last *m*-mer in the new *k*-mer for a faster calculation of the current *k*-mer’s minimizer).

## 4 Results

We tested the performance of AdaOrder and several popular orders on multiple datasets, and also compared the performance of DGerbil (which uses AdaOrder), FGerbil (which uses frequency), and the original Gerbil.

### Algorithm 3 DGerbil

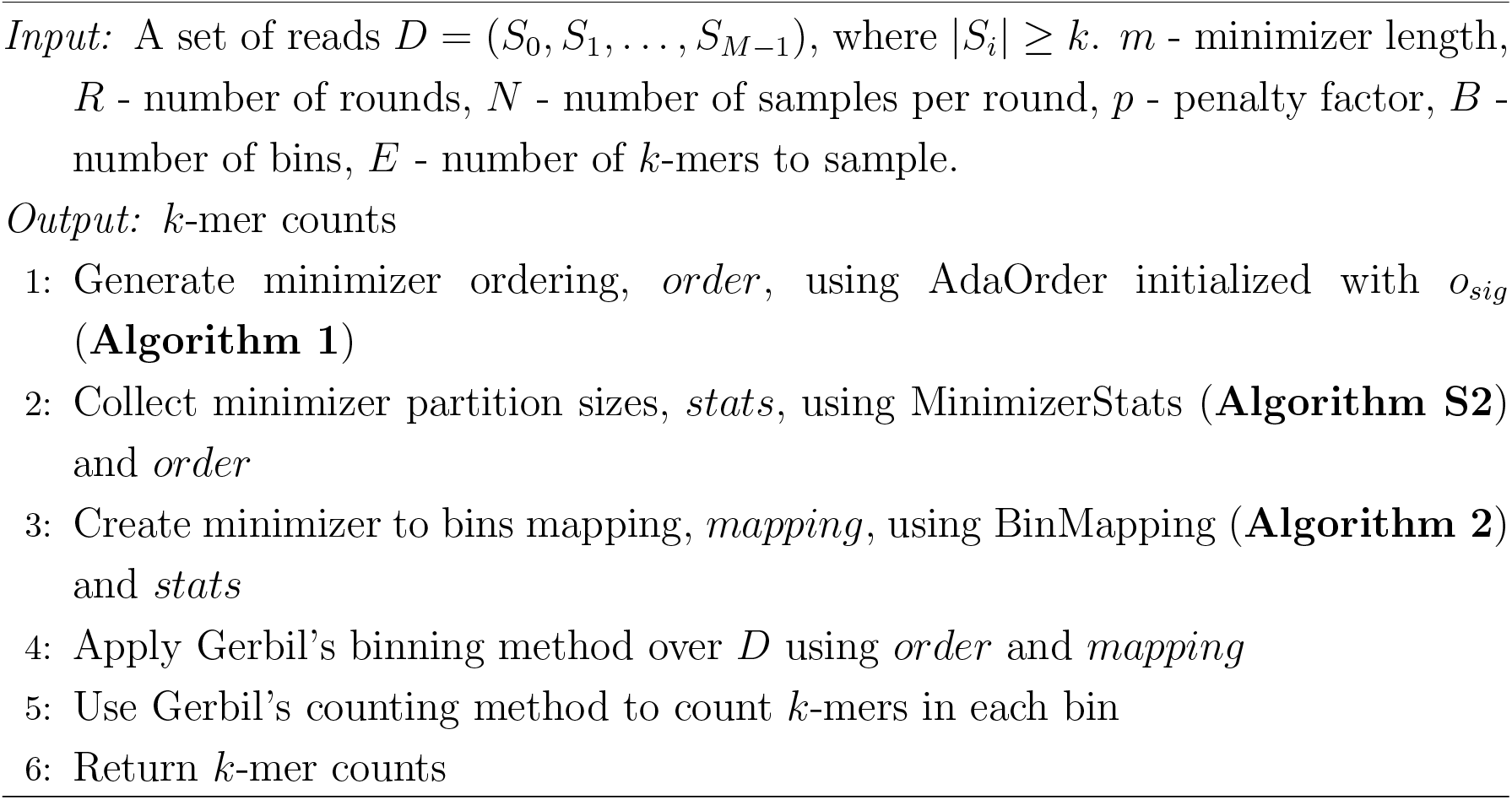

We used eight datasets; seven were those used in a recent *k*-mer counter benchmark [13] and the eighth was a large freshwater metagenomic dataset (FW [15], SRR6787039). See **Table 1** for the properties of the datasets. Both AT and NC are long reads datasets collected using a PacBio technology.

**Table 1:** Characteristics of the benchmark datasets. Note that the SRA toolkit merges pair-end reads into a single read by default when downloading, and therefore the pair-end reads of HS2, MB, DM, FV and FW were merged and the reported read length is after the merge.

| Dataset | Size ($10^9$ bytes) | Avg. read length | Species |
| --- | --- | --- | --- |
| HS1 | 292 | 151 | H. sapiens |
| HS2 | 347 | 202 | H. sapiens |
| AT | 72.7 | 4804 | A. Thaliana |
| FW | 406 | 300 | Fresh water metagenome |
| FV | 14.1 | 508 | F. vesca |
| NC | 45.9 | 7778 | N. crassa |
| DM | 10.7 | 152 | D. melanogaster |
| MB | 198 | 101 | M. balbisiana |

Throughout this section, the default parameters used for AdaOrder unless stated otherwise were *o*_*init*_ = *o*_*sig*_, *R* = 10^4^, *N* = 10^5^, and *p* = 0.01. The choice of these parameters is discussed in **Supplementary Section S3**. For the frequency order we collected statistics over the entire dataset. Minimizer length was always *m* = 7.

### 4.1 AdaOrder reduces maximum load and unevenness

We applied the lexicographic, signature, random, frequency and AdaOrder orders on four large datasets for *k* = 28 and 55. In the random order we used a different random mask for each generated random order. Maximum load and unevenness results are shown in **Table 2** and **Table 3**, respectively. Frequency order consistently had the lowest maximum load and unevenness followed by AdaOrder, while the predefined orders were substantially worse.

**Table 2:** Maximum load ×10^−6^.

| Dataset | $k$ | Lexicographic | Random | Signature | AdaOrder | Frequency |
| --- | --- | --- | --- | --- | --- | --- |
| HS1 | 28 | 78.5 | 20.1 | 14.6 | 5.7 | 3.0 |
|  | 55 | 157.1 | 51.7 | 36.7 | 10.2 | 4.8 |
| HS2 | 28 | 83.7 | 22.1 | 15.6 | 5.8 | 3.1 |
|  | 55 | 175.3 | 60.0 | 41.7 | 9.9 | 5.1 |
| FW | 28 | 58.1 | 54.0 | 27.1 | 11.0 | 5.3 |
|  | 55 | 112.3 | 70.2 | 58.8 | 19.0 | 8.6 |
| AT | 28 | 35.1 | 4.7 | 12.0 | 0.6 | 0.5 |
|  | 55 | 6.0 | 1.2 | 3.7 | 0.8 | 0.3 |

**Table 3:** Unevenness ×10^7^.

| Dataset | $k$ | Lexicographic | Random | Signature | AdaOrder | Frequency |
| --- | --- | --- | --- | --- | --- | --- |
| HS1 | 28 | 4.6 | 2.6 | 2.5 | 1.0 | 0.4 |
|  | 55 | 11.4 | 4.5 | 5.2 | 1.6 | 0.6 |
| HS2 | 28 | 4.6 | 2.0 | 2.5 | 1.0 | 0.4 |
|  | 55 | 11.6 | 3.7 | 5.2 | 1.5 | 0.5 |
| FW | 28 | 1.8 | 2.1 | 1.8 | 0.9 | 0.3 |
|  | 55 | 4.2 | 4.4 | 3.8 | 1.9 | 0.6 |
| AT | 28 | 18.0 | 3.1 | 4.0 | 0.7 | 0.2 |
|  | 55 | 168.5 | 15.9 | 63.4 | 5.7 | 2.2 |

We also calculated the distribution of load across minimizers for each order. In **Figure 2a** we plot the load of the 1000 minimizers with the highest load on HS2 using *k* = 55 for the different orders. AdaOrder and frequency distributed the *k*-mers much more evenly across these minimizers than the other orders, in line with the results in **Table 2** and **Table 3. Figure 2b** shows the cumulative distribution of the load for all minimizers. AdaOrder and frequency order did a good job in balancing the top loads compared to the other orders. For example, when using signature, ∼ 20% of *k*-mers were covered by the top 20 minimizers, while AdaOrder used the 100 top minimizers to cover ∼ 20% of the *k*-mers. The *number* of minimizers used also differed substantially between orders: 7731 for frequency, 4940 for AdaOrder, 4939 for signature, 2694 for lexicographic and 3828 for the random order. Interestingly, AdaOrder achieved much lower and more even maximum loads than the signature order while still using a very similar number of minimizer partitions, whereas frequency order used many more minimizers to achieve better performance.

**Figure 2:**
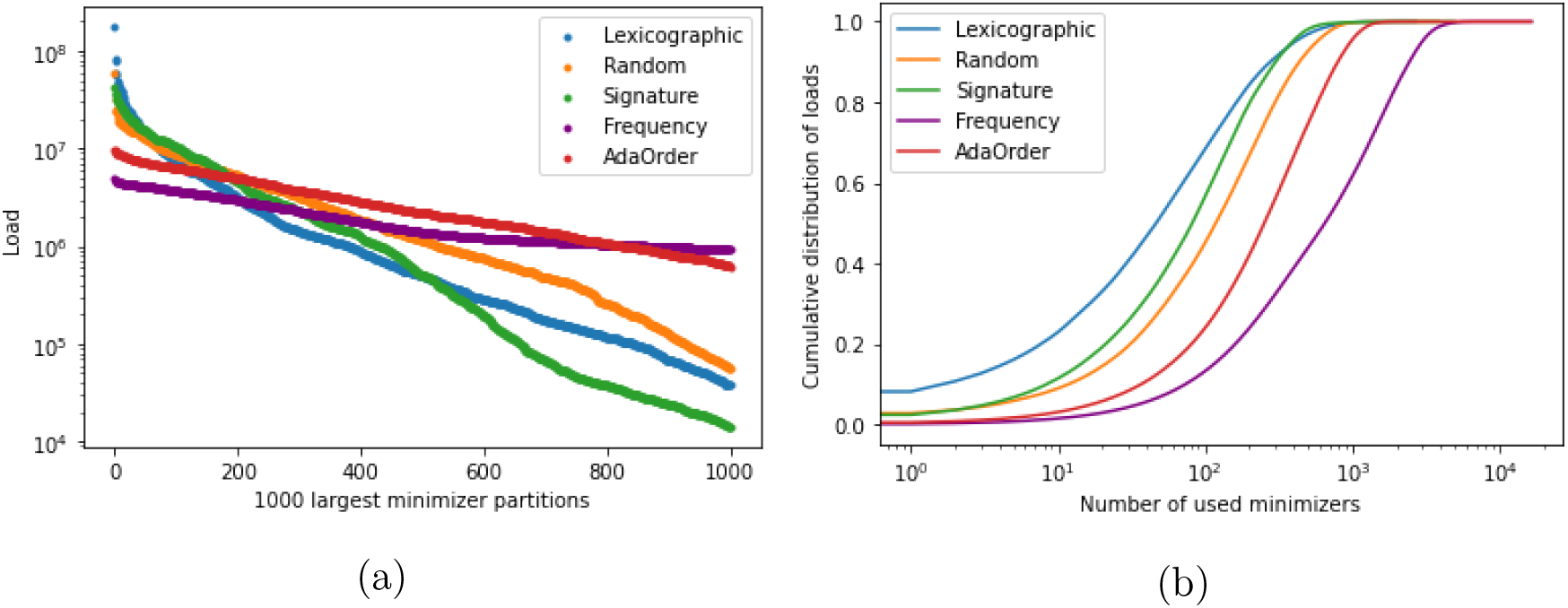
Distribution of loads across minimizers. Results are for HS2 with *k* = 55. (a) Load of the 1000 minimizers with highest load. (b) Cumulative distribution of the load of all minimizers. In both figures minimizers were sorted in decreasing load for each order.

### 4.2 The effect of *R, N* and *p* on maximum load and unevenness

We explored the impact of each of the input parameters *R, N*, and *p* of AdaOrder on the maximum load and unevenness. This analysis can help in the choice of parameters in AdaOrder. In **Supplementary Section S1** we performed a statistical analysis of the accuracy of sampling-based load estimation, which sheds light on the recommended size of *N*, We tested various combinations of *R, N* and *p*. In each test we fixed two parameters to their default values and varied the third. All the tests were performed on the HS2 dataset with *k* = 55.

**Figure 3a** shows the effect of varying *R*. We observe that both maximum load and unevenness drop drastically from the baseline within relatively few rounds (note that the x axis is logarithmic). The load plateaus after around 10^4^ rounds, while unevenness keeps decreasing slowly.

**Figure 3:**
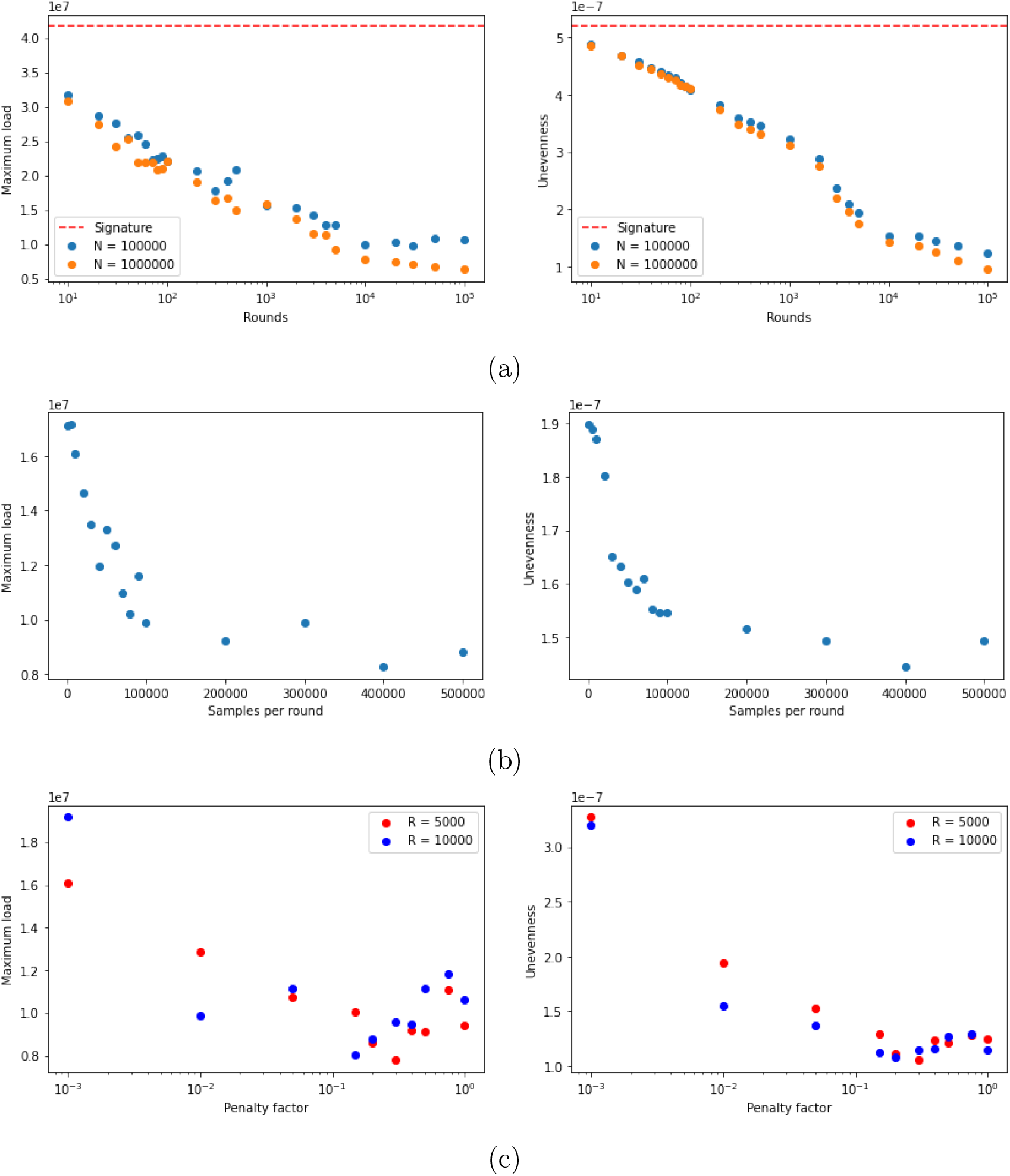
Maximum load and unevenness when varying parameters *R, N* and *p* in AdaOrder. Each graph shows the performance when varying a single parameter. (a) Impact of varying *R*, the number of rounds. The dashed red line corresponds to the maximum load and unevenness value of the signature order with which it is initialized. (b) Impact of changing the sample size *N*. (c) Impact of the penalty factor *p*.

**Figure 3b** shows the effect of varying *N*. Both maximum load and unevenness decrease as *N* gets larger. This demonstrates the benefit of choosing a large sample.

**Figure 3c** shows the effect of varying *p*. The load decreases first as *p* increases, but no clear trend is observed for *p* ≥ 0.1, with substantial jumps in the load. A similar trend but with lower variability is observed for unevenness. The trends are similar when using 5000 and 10000 rounds. The high variability for high values of *p* is likely due to large changes in the order in the final rounds after all of the top loads have already been made relatively even. **Supplementary Figure S1** shows a similar analysis for *N* = 10^4^ − 10^8^ and different rounds.

Based on the analysis above and the theoretical analysis in **Supplementary Section S1** we established recommended parameters for running AdaOrder. See **Supplementary Section S3** for details.

### 4.3 Transfering the order across datasets

We wished to see whether an order produced on one dataset can assist in creating orders for other datasets from the same species, either by running it as is, or by using it as a starting order in the optimization for the other datasets. We reasoned that if this is the case, then it could speed up the process of producing good orders for other datasets from the same species.

Suppose we have a source dataset *src* and a destination dataset *dst*. We apply AdaOrder on *src* and use that order on *dst*. We performed two experiments: One where *src* was HS1 and *dst* was HS2, and another where *src* was HS2 and *dst* was HS1. We ran AdaOrder with *k* = 28 and *k* = 55 over *src* and tested the efficiency of the minimizer order it produced over *dst*. We also ran AdaOrder over *dst*, initialized with the order produced by AdaOrder for *src*.

The results are shown in **Figure 4** and **Figure 5**. We see that transferring the order works similarly to the AdaOrder results produced for the destination dataset independently of *src* (the dashed green line). Moreover, starting from the *src* order, the algorithm can improve unevenness and maximum load within far fewer optimization rounds compared to running AdaOrder for *dst* from scratch (*<* 1000 vs. 10^4^).

**Figure 4:**
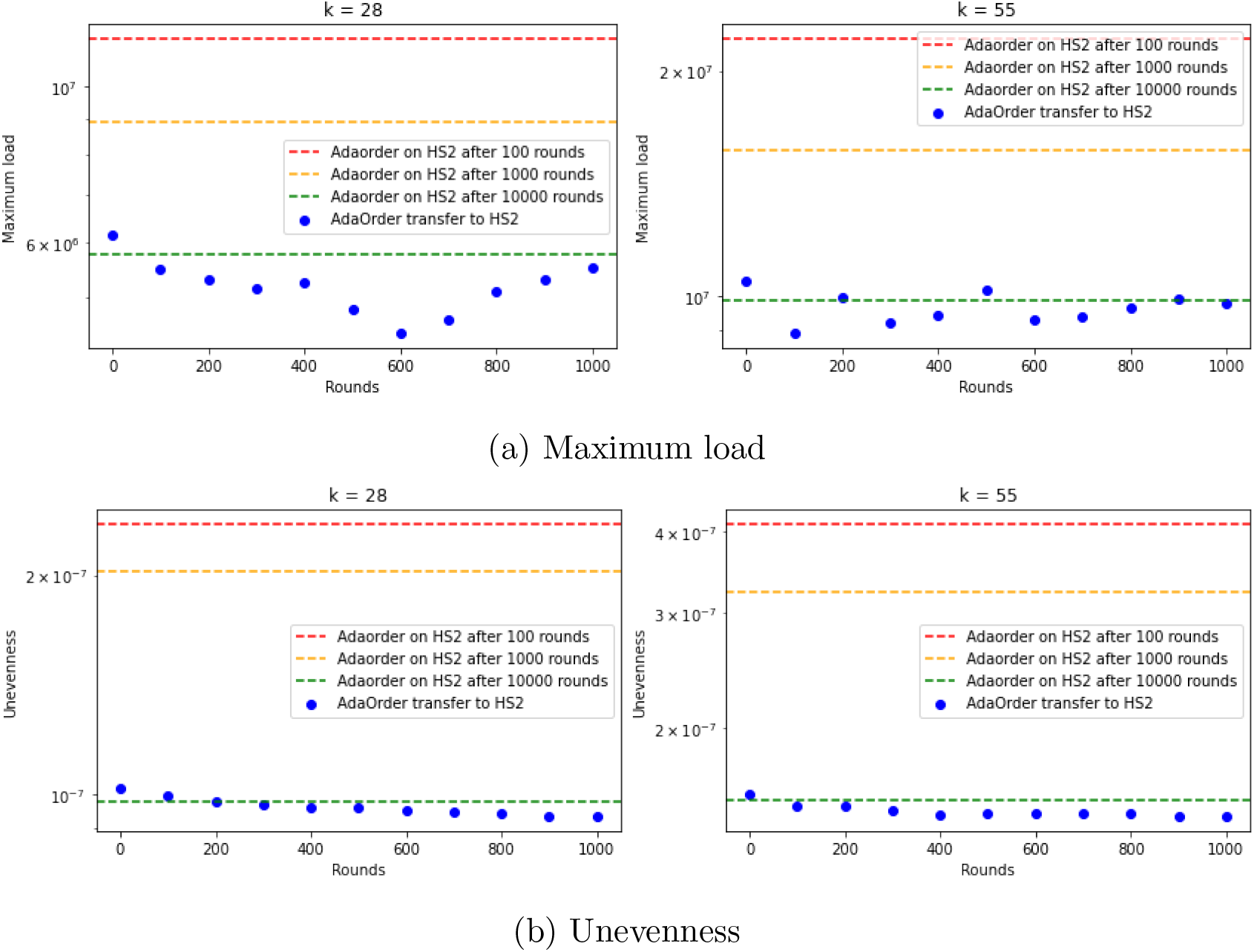
Order transfer from HS1 to HS2. The value at round 0 corresponds to applying the HS1 order as is on HS2. The dashed lines correspond to the values obtained by running AdaOrder on HS2 with the default initialization, for different number of rounds.

**Figure 5:**
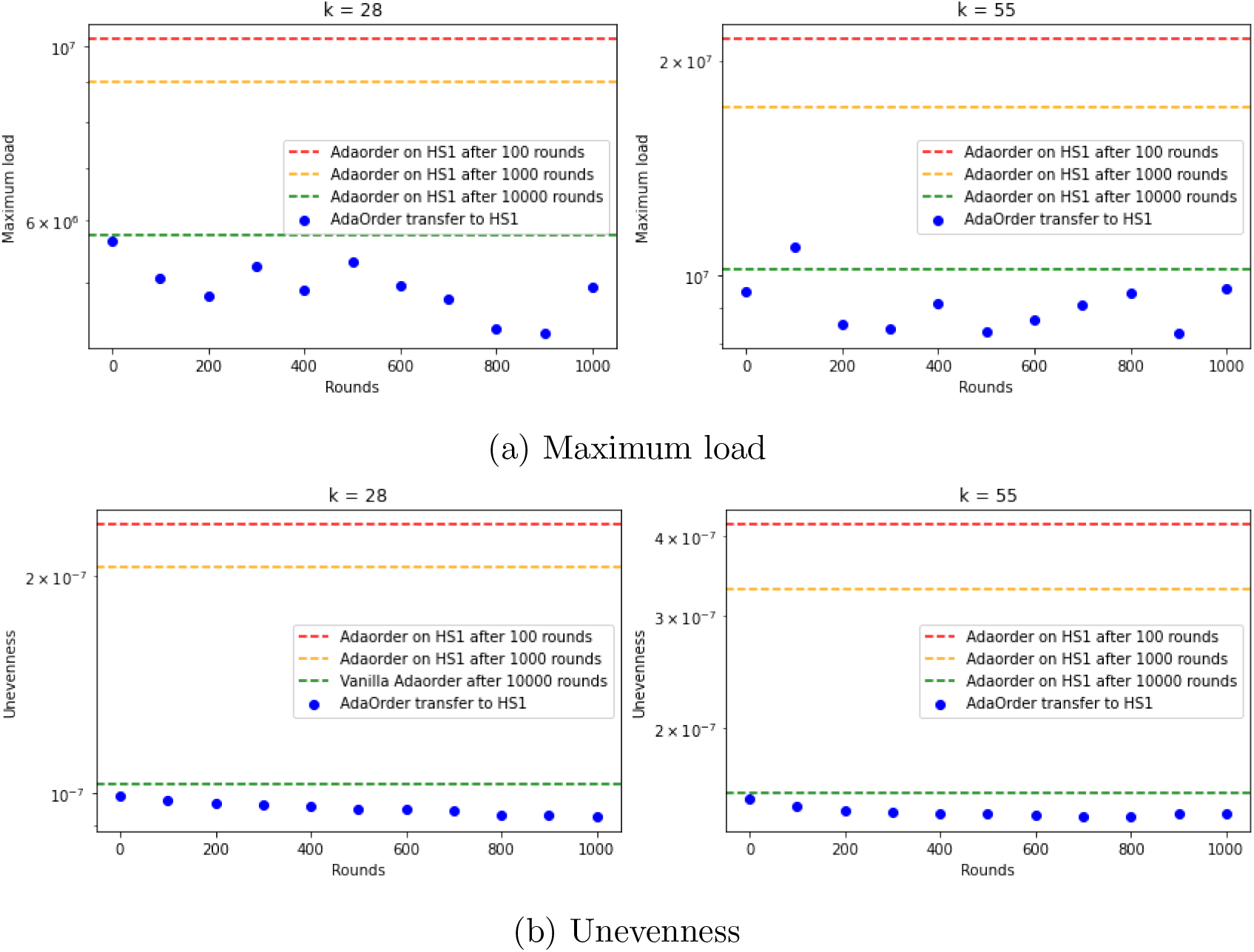
Order transfer from HS2 to HS1. The value at round 0 corresponds to applying the HS2 order as is. The dashed lines correspond to the values obtained by running AdaOrder on HS1 with the default initialization, for different number of rounds.

We conclude that the order can be transferred across datasets from the same species, saving most of the running time and preserving the order quality.

### 4.4 DGerbil reduces memory usage

We compared the performance of Gerbil, DGerbil and FGerbil. All algorithms were run with 512 bins and *k* = 28, 55, 70, 90 on each dataset (28 and 55 were used in [13]). All the experiments were measured on a 128-core server with 64 3.35 GHz CPUs and 1000GB of RAM (AMD EPYC 7702). We ran all algorithms with 12 threads, the same number of threads as in [13]. The storage used in our experiments is an external NFS storage, and all IO operations were performed against it (e.g., reading datasets and writing temporary and output files). The RAM usage of AdaOrder was externally limited to a maximum of 1.6GB (we used Java’s Xmx flag to limit the program’s heap size).

The results for the four datasets with the highest RAM usage are shown in **Figure 6**. The results for the other four datasets are in **Supplementary Figure S2**. The figures show the runtime and RAM usage on each dataset. For DGerbil (respectively, FGerbil) the time of AdaOrder (respectively, frequency order) is shown separately. DGerbil used consistently lower RAM except when using the smallest value of *k* (28). All tools tended to use more memory as *k* increased. Similar RAM trends were observed for the other datasets (**Supplementary Figure S2**) though absolute numbers were lower.

**Figure 6:**
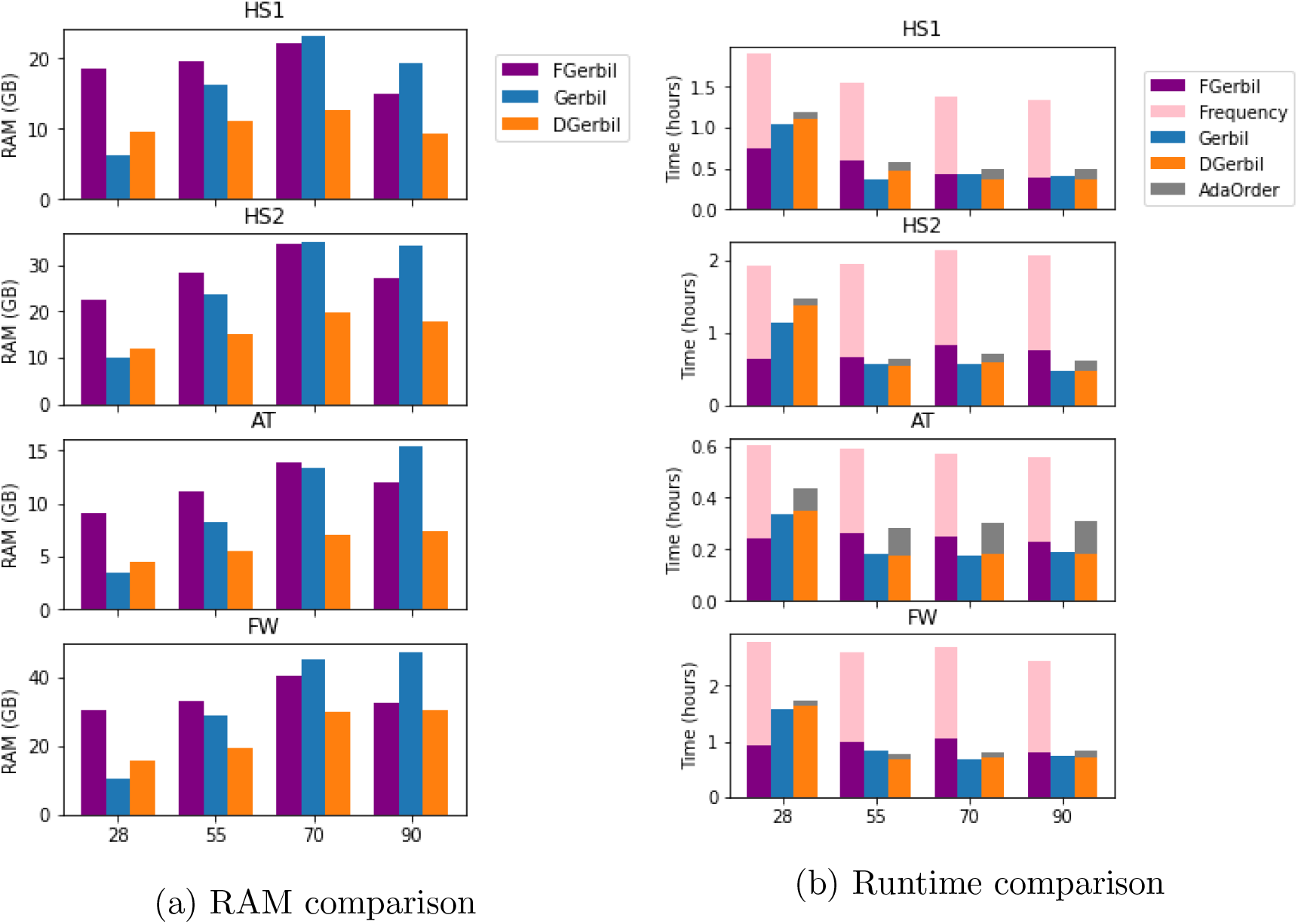
Performance of Gerbil, FGerbil and DGerbil. In (b), for DGerbil and FGerbil, AdaOrder and Frequency times for creating the order and collecting binning statistics are shown separately.

In terms of time, it took an average of 6m 50s to run AdaOrder and to collect statistics for binning. This runtime was roughly constant as the sampling process does not depend on the dataset size. For frequency it took on average about 39 minutes to finish collecting statistics for the order and for the binning. For the smaller datasets shown in **Supplementary Figure S2** the runtime of Gerbil was already very low and therefore the time to run AdaOrder or frequency dominated that of the *k*-mer counting.

## 5 Discussion

We developed AdaOrder, an algorithm for creating an improved minimizer order on a given dataset by repeatedly adjusting the order to reduce the maximum load. AdaOrder significantly reduced maximum load and unevenness compared to the signature order and to other predefined orders. The performance of the frequency order was even better, but it did not translate to a better *k*-mer counting algorithm (see below). AdaOrder was integrated into the *k*-mer counter Gerbil together with a *k*-mer sampling method in order to map minimizers into bins. This reduced memory usage by 30-50% of Gerbil for large *k*.

AdaOrder has a roughly constant runtime of under seven minutes regardless of the size of the dataset. For small datasets the absolute reduction in the memory usage is minor, and the running time of AdaOrder dominates the *k*-mer counting process, so using AdaOrder is not advantageous. The main advantage of AdaOrder is on larger datasets, especially when longer *k*-mers are used. In these cases the memory savings are substantial, while the additional time required to run AdaOrder is relatively minor.

DGerbil outperformed Gerbil in memory, but required a bit more time. By transferring an order precomputed by AdaOrder on another dataset of the same species, most or all of this time can be saved, thus matching Gerbil’s time and cutting memory by 30% − 50% on large *k* values.

Interestingly, applying only one of our modifications to Gerbil individually – using AdaOrder instead of signature order, or mapping minimizers to bins based on *k*-mer sampling statistics – actually *increased* the memory usage (results not shown). Hence, we suspect that multiple factors (maximum load, the algorithm for bin mapping and details of the implementation) together determine the actual memory usage of Gerbil.

Specific implementation details of Gerbil may also explain some of the other results we observed. For example, Gerbil decides in advance what hash table size to use for a bin based on the number of *k*-mers written to it, and on the ratio of distinct *k*-mers to total *k*-mers in the previous bin. Choosing a hash table size that is too big wastes memory, while choosing one that is too small increases running time. Since this heuristic and its parameters were optimized in Gerbil, this may explain why Gerbil has better performance for the smallest *k* values tested (*k* = 28). Similar considerations may explain why the frequency order fails to improve Gerbil: The distribution of minimizer loads is very different from what Gerbil expects and is implemented to optimize.

Several areas for improvement and open questions remain. Most importantly, can one improve the algorithm by dynamically choosing values for the parameters *R, N*, and *p*? (i) Our analysis suggests that the number of samples per round should not be fixed throughout the entire process, as it depends both on the maximum relative load and on the difference between the top relative loads, both of which decrease as the algorithm progresses. (ii) Having a stopping condition for the entire process instead of a fixed number of rounds could be beneficial, as we observed that AdaOrder continues to alter the order even after it it is no longer beneficial. (iii) One may want to penalize the maximum load minimizer in a round as a function of how large the estimated maximum relative load is (i.e., if the maximum load is larger then we would like *p* to be larger). Together, some or all of these ideas could improve the algorithm further.

The frequency order achieved superior maximum load and unevenness results compared to AdaOrder. However, its use in Gerbil required more RAM compared to DGerbil. An interesting topic for future research is the design of a *k*-mer counter with low memory usage based on the frequency order. Leveraging frequency’s potential in this task may lead to an even more memory efficient and faster *k*-mer counter.

We have demonstrated the utility of directly optimizing the maximum load of a minimizer order in binning applications. Using AdaOrder in Gerbil further improved its RAM usage, especially for longer *k*-mers. This approach has potential to reduce the memory footprint of other sequence analysis algorithms on large datasets.

## 6 Acknowledgments

We thank Nimrod Rappoport and Jonathan Schifter for helpful advice. Study supported in part by the Israel Science Foundation (grant No. 3165/19, within the Israel Precision Medicine Partnership program, and grant No. 1339/18), by Len Blavatnik and the Blavatnik Family foundation, and by the Zimin Institute for Engineering Solutions Advancing Better Lives.

## Supplementary information

### S1 Accuracy of sampling-based load estimation

A key parameter in AdaOrder is *N*, the number of *k*-mers it samples in each round. If *N* is too small, then the penalized minimizer may not actually have a true high load. If it samples too many *k*-mers then the output minimizer scheme may be superior, but RAM usage and running time will increase. Here we provide a theoretical analysis to guide the choice of *N* so as to balance AdaOrder’s resource usage and the performance of the output minimizer scheme.

We bound the probability of deviation of an estimated load from its true value as a function of *N* as follows. Let *x* be a minimizer with true relative load *r* on dataset *D*. Let *S* ⊆ *D* be a subset with *N* sampled *k*-mers. Since *k* is long and *N* is relatively small compared to the number of possible *k*-mers, we make the assumption that the sampled *k*-mers are distinct. Assume that the sampling process is binomial, namely, *x* has probability *r* to be the minimizer of a *k*-mer. Then for a large enough *N* we expect *x*’s sample load to be

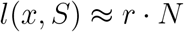

We are interested in choosing *N* so that the difference from the true relative load is less than *δ*. The probability of that event is

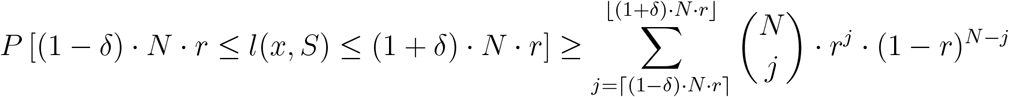

The expression on the left describes the chance that a sampled load differs by at most a factor of *δ* from the expected sampled load. We lower-bounded this probability with the binomial sum on the right, denoted as *ρ*(*N, r, δ*). This bound can guide us in choosing a value for *N*, by computing it for typical *r* values, and a small enough value for *δ*.

Our empirical results showed that for *m* = 7, maximum load *m*-mers have relative load of ∼ 0.01 − 0.06 for pre-defined orders used in practice (e.g., signature or lexicographic). For example, in one human dataset (HS2, see below) with *k* = 55 the maximum load was 0.056 for lexicographic and 0.013 for signature. **Table S1** provides values for *ρ*(*N, r, δ*) using different combinations of *N* and *r*, with a fixed *δ* = 0.1. We see that *N* = 50000 − 100000 gives a high probability to estimate the load of *m*-mers accurately for realistic loads. (For *m >* 7 we expect the maximum relative load to be smaller, requiring larger values of *N*.)

**Table S1:** Sampling estimation accuracy for different values *N, r* and for *δ* = 0.1.

|  |  |  |  |  |  |
| --- | --- | --- | --- | --- | --- |
| $r$ | 0.05 | 0.01 | | 0.002 | |
| $N$ | 10000 | 10000 | 50000 | 50000 | 100000 |
| $\rho(N, r, \delta)$ | 0.97 | 0.68 | 0.97 | 0.68 | 0.84 |

The above bound is for the load estimate of a single minimizer, but is not necessarily accurate for the *maximum* load. How good the estimate of the maximum load minimizer is depends on how close the second highest load is: if it is close to the maximum, then the identified maximum load minimizer may be incorrect, while if it is far from the maximum, then we can likely estimate the correct maximum minimizer with high accuracy.

In AdaOrder, by design, the top loads decrease and get closer to each other as the iterations progress. We therefore want to have a low probability for a minimizer with a small relative load to end up with the largest count when sampling in an iteration. We do not necessarily expect AdaOrder to detect the exact minimizer with the highest load, but rather a minimizer with a high relative load (e.g., with one of the *n* highest relative loads). Therefore we wish to bound the probability that a minimizer with a low true relative load 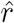 that is smaller than the top *n* relative loads will mistakenly have an estimated sample load *X* larger than all of the top *n* loads.

Assume that the relative loads for the top *n* minimizers are *r*_1_ ≥ *r*_2_ ≥ … ≥ *r*_*n*_, with corresponding sample loads *X*_1_, …, *X*_*n*_. Assuming the estimates are independent (since *N* is very large), we get:

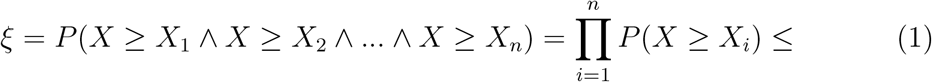

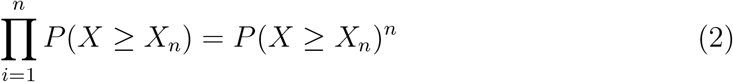

We evaluated this expression for *n* = 10, using the loads observed in the last round of AdaOrder in the HS2 dataset with *k* = 55: The maximum load was 0.00318, and the 10th largest load was 0.00279. Suppose 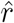 equals 80% of the maximum relative load in our example (0.00254). We empirically evaluated *P* (*X* ≥ *X*_*n*_) by sampling 10^5^ times from the binomial distribution with parameters 0.00279 and *N* = 10^5^ trials, obtaining *P* (*X* ≥ *X*_*n*_) = 0.136 and

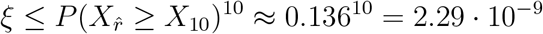

This calculation suggests that for *N* = 10^5^ there is a very low chance of a gross mistake in the identification of a minimizer with a high load.

In **Supplementary Section S2** we show the effect of varying the number of samples, *N*, in different rounds of AdaOrder, demonstrating the effect described.

### S2 The Effect of sampling depth on accuracy of load estimation

We tested how the sampling depth and the number of rounds affect AdaOrder. We used dataset HS2 with *p* = 0.01. **Supplementary Figure S1** shows the relative load as a function of the sample size, for different rounds of the order optimization process. Namely, for each *N*, the minimizer that would have been selected if sampling stopped after *N k*-mers, and its relative load are shown. Results are shown for the first, 1000th and 10^4^th round. We see that in the first round the identity and relative load of the maximum load minimizer could be estimated with only 2 · 10^4^ samples. However, in later rounds, as the loads get more even, the load of the chosen minimizer improves even after 10^5^ samples, and it takes even more samples for the identity of the minimizer to stabilize.

### S3 Recommended parameters for AdaOrder

We selected the default parameters of *R* = 10^4^, *N* = 10^5^, *p* = 0.01 for AdaOrder. These parameters were chosen to limit the runtime of AdaOrder while still achieving a good ordering with low maximum loads (and potentially low RAM usage in DGerbil).

According to **Figure 3a**, increasing *R* above 10^4^ had little effect on the maximum load. The same figure also suggests that *N* = 10^5^ did not have a notably worse maximum load than *N* = 10^6^. Lower values of *N* performed worse, as shown in **Figures 3b** and **S1**. In fact, **Supplementary Figure S1** seems to indicate that more samples are needed in the later rounds compared to the initial rounds, since the maximum relative load is getting smaller as AdaOrder progresses (see **Supplementary Table S1**). *N* = 10^5^ thus gives a good balance between runtime and maximum load.

The penalty factor *p* = 0.01 is a conservative choice. Values of 0.1 and above gave unstable load and unevenness. Alternatively, one can choose 0.01 *< p* ≤ 0.1, in combination with lower *R*, as **Figure 3c** suggests that somewhat higher *p* can still lead to more efficient flattening of the order, and thus require fewer rounds.

**Figure S1:**
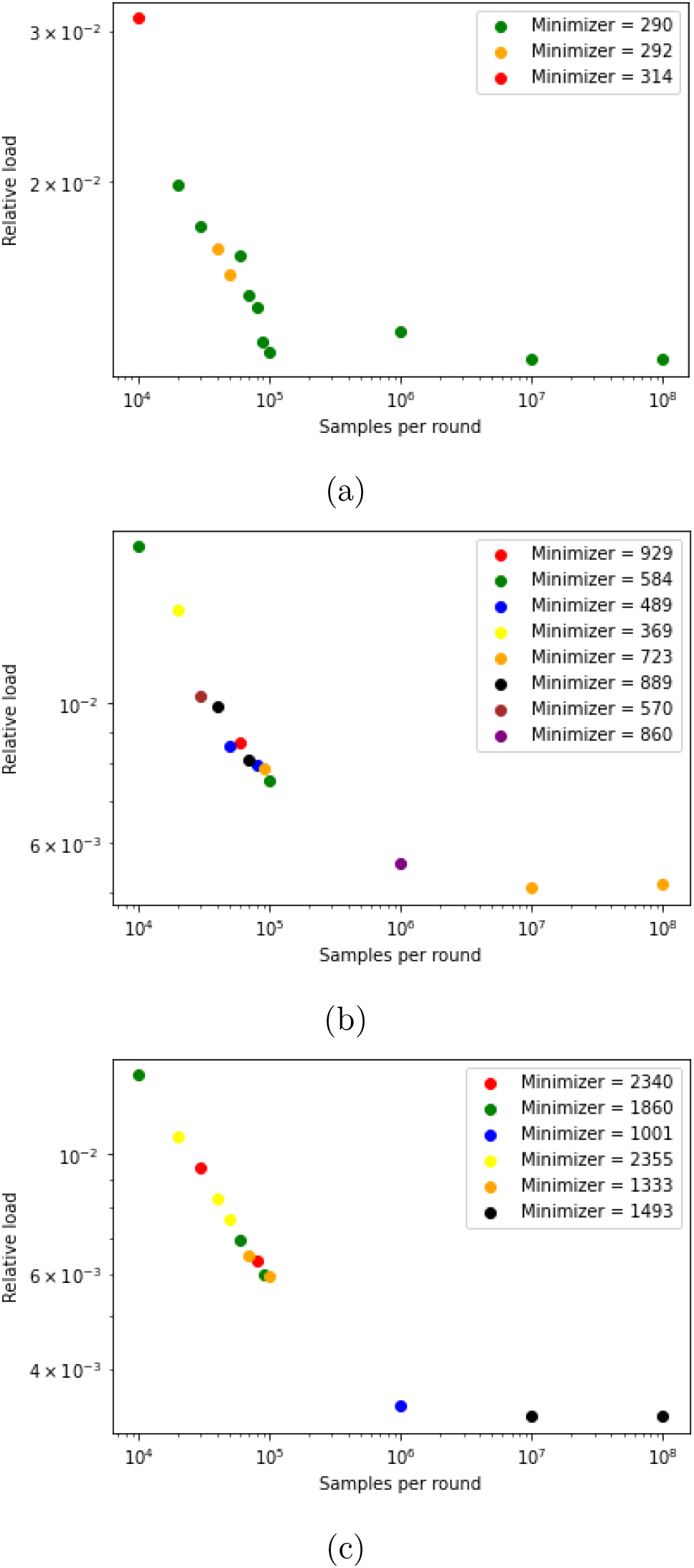
Effect of the sample size and round number on the estimate of maximum load minimizer. (a) 1st round. (b) 1000th round. (c) 10^4^th round. Each dot shows the minimizer to be selected based on the corresponding sample size. The y-axis is the relative load of the selected minimizer. The numbers assigned as minimizer labels are irrelevant, but the same color indicates a repeated minimizer.

#### Algorithm S1 KMC3 mapping outline

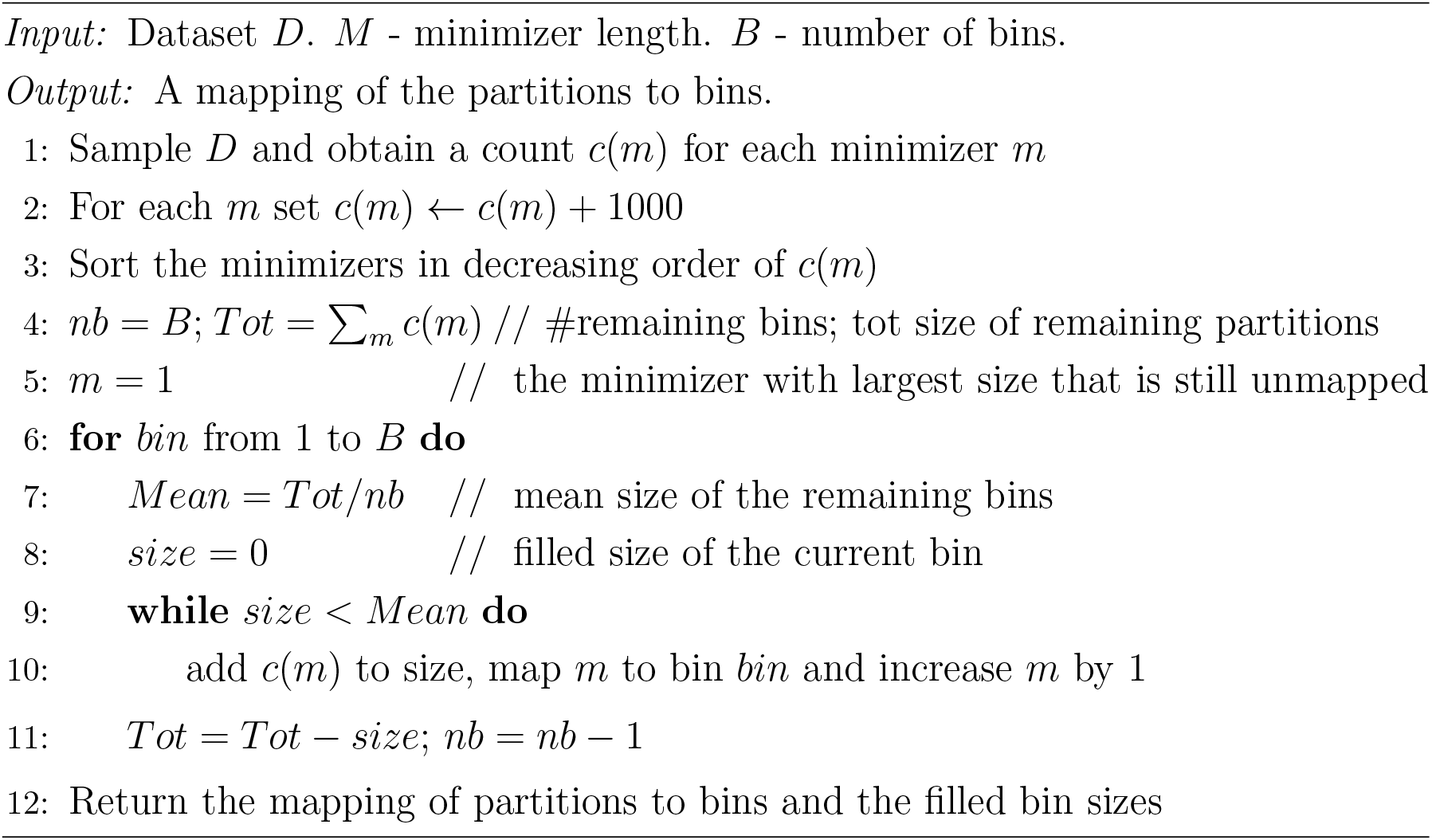

#### Algorithm S2 MinimizerStats

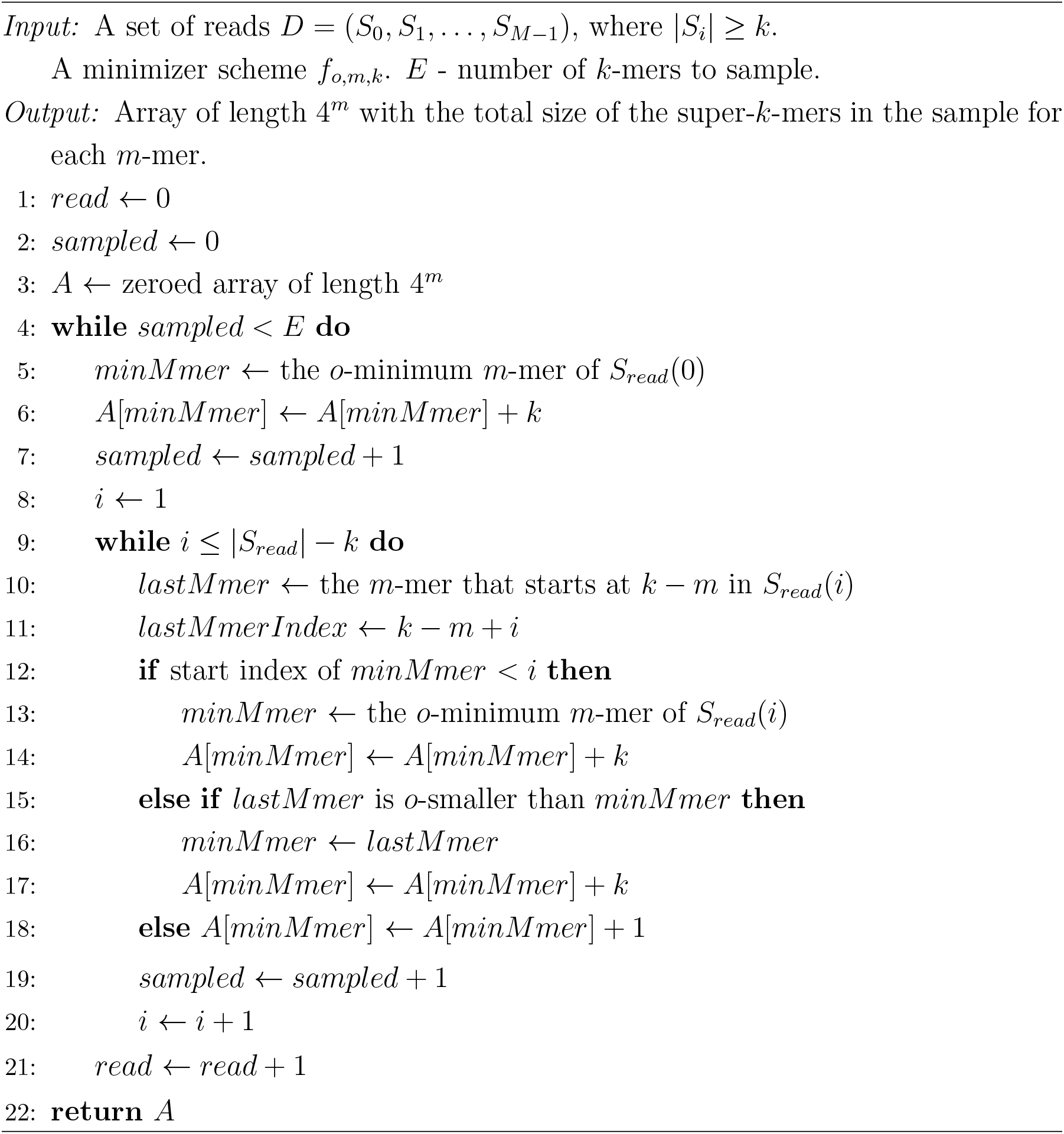

**Figure S2:**
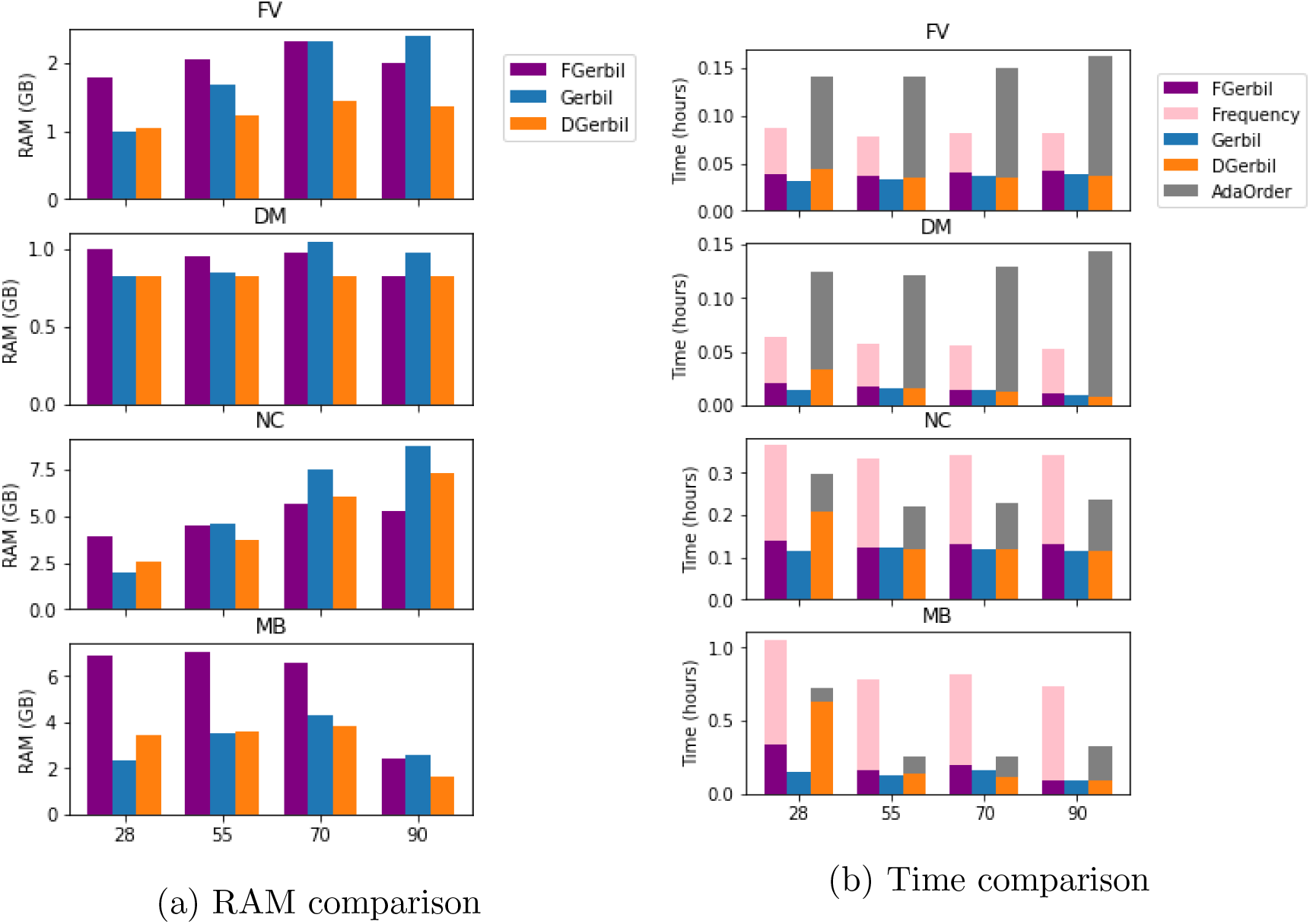
Performance of Gerbil, FGerbil and DGerbil. In (b), for DGerbil and FGerbil, AdaOrder and Frequency times for creating the order and collecting binning statistics are shown separately.

